# Dissociated amplitude and phase effects of alpha oscillation in a nested structure of rhythm- and sequence-based temporal expectation

**DOI:** 10.1101/2022.09.01.506156

**Authors:** Zhongbin Su, Xiaolin Zhou, Lihui Wang

**Affiliations:** Shanghai Key Laboratory of Psychotic Disorders, Shanghai Mental Health Center, Shanghai Jiao Tong University School of Medicine, Shanghai, China; Institute of Psychology and Behavioral Science, Shanghai Jiao Tong University, Shanghai, China; Beijing Key Laboratory of Behavior and Mental Health, School of Psychological and Cognitive Sciences, Peking University, Beijing, China; Shanghai Key Laboratory of Mental Health and Psychological Crisis Intervention, School of Psychology and Cognitive Science, East China Normal University, Shanghai, China; PKU-IDG/McGovern Institute for Brain Research, Peking University, Beijing, China; Shanghai Center for Brain Science and Brain-Inspired Intelligence Technology, Shanghai, China

**Keywords:** temporal expectation, neural entrainment, pre-stimulus oscillation, alpha amplitude, alpha phase

## Abstract

The human brain can utilize various information to form temporal expectation and optimize perceptual performance. Here we show dissociated amplitude and phase effects of pre-stimulus alpha oscillation in a nested structure of rhythm- and sequence-based expectation. A visual stream of rhythmic stimuli was presented in a fixed sequence such that their temporal positions could be predicted by either the low-frequency rhythm, the sequence, or the combination. The behavioral modelling indicated that rhythmic and sequence information additively led to increased accumulation of sensory evidence and alleviated threshold for the perceptual discrimination of the expected stimulus. The electroencephalographical (EEG) results showed that the alpha amplitude was dominated by rhythmic information, with the amplitude fluctuating in the same frequency of the oscillation entrained by the rhythmic information (i.e., phase-amplitude coupling). The alpha phase, however, was affected by both rhythmic and sequence information. Importantly, rhythm-based expectation improved the perceptual performance by decreasing the alpha amplitude, whereas sequence-based expectation did not further decrease the amplitude on top of rhythm-based expectation. Moreover, rhythm-based and sequence-based expectation collaboratively improved the perceptual performance by biasing the alpha oscillation toward the optimal phase. Our findings suggested flexible coordination of multiscale brain oscillations in dealing with a complex environment.

## Introduction

We live in a world teemed with temporal regularities: day and night alternate with each other approximately every 12 hours; the thunder is always heard after seeing the lightning. Such temporal regularity affords the temporal expectation of an event, which avails to guide perception and action that serve the current goal (Jones, 1976). A few minutes after leaving home for work in a morning routine, for instance, one can quickly spot the cafe and grab a coffee without paying attention to the surroundings. In laboratory settings, a stimuli stream with temporal regularity can lead to improved task performance such as increased accuracy and/or facilitated reaction times (RTs) in discriminating the perceptual attributes of the stimulus embedded in the stream (Cravo, Rohenkohl, Wyart, & Nobre, 2013; Morillon, Schroeder, Wyart, & Arnal, 2016; Sanabria, Capizzi, & Correa, 2011). The improved perceptual performance by expectation was supported by sharpened neural representation (De Lange, Heilbron, & Kok, 2018) and earlier neural excitation (Anderson & Sheinberg, 2008) for the expected event.

Temporal expectation can be built upon multiple bases. In a flow of stimuli, both the rhythmic and the sequence regularity of the stimuli can afford the temporal expectation of a particular stimulus (Nobre & Van Ede, 2018). For instance, a red circle appears every 800 ms (i.e., a regular rhythm of 1.25 Hz), or always comes after two successive blue circles (i.e., a regular sequence). In the case of a regular sequence, the successive stimuli were not necessarily presented in a certain rhythm, and hence sequence-based temporal expectation is dissociable from the rhythm-based expectation. Although either the rhythmic or the sequence regularity alone can be predictive of a stimulus, and there is already evidence suggesting both common and dissociable mechanisms of different forms of temporal expectation (Breska & Deouell, 2017; Correa, Cona, Arbula, Vallesi, & Bisiacchi, 2014), it remains unknown how the nested rhythmic and sequence information are combined to affect the sensorimotor processing and the corresponding behavioural performance. This is of significance as different sources of temporal regularity are often nested in a real-world situation.

The rhythmic regularity has been associated with neural entrainment, a process where the intrinsic neural oscillations are tuned to the rhythm of the external stimuli (Calderone, Lakatos, Butler, & Castellanos, 2014; Haegens & Golumbic, 2018; Jones, 2010). Such entrainment can be maintained even for a few cycles after the offset of the external rhythm, and meanwhile can still influence behavioral performance (Lakatos et al., 2013; Spaak, De Lange, & Jensen, 2014). It has been consistently shown in various contexts (e.g., visual, auditory, multisensory) that the perceptual performance was improved after the neural signals were entrained by the rhythmic stimuli (De Graaf et al., 2013; Lakatos, Karmos, Mehta, Ulbert, & Schroeder, 2008; Mathewson et al., 2012). By entrainment, a stimulus gains optimized processing when aligned with the peak of the oscillatory neural signals (i.e., high-excitability phase) relative to the trough of the oscillatory signals (i.e., low-excitability phase), and the perceptual performance was critically mediated by the phase concentration of the entrained activity (Cravo et al., 2013; Stefanics et al., 2010). It has been shown that the phase concentration of the low-frequency (i.e., delta-band) oscillation not only underlined the temporal expectation afforded by the rhythmic stimuli, but also the temporal expectation based on memory (Breska & Deouell, 2017), suggesting a general role of phase concentration in multiple forms of temporal expectation.

Another important component of brain activity that has been linked to temporal expectation is the alpha-band (around 8-12Hz) oscillation, which is suggested as causal for the timing and perceptual processing of the upcoming stimulus (Klimesch, 2012; Romei, Gross, & Thut, 2010). On the one hand, the pre-stimulus alpha oscillation showed decreased amplitude during temporal expectation, indicating increased neural excitation in preparation for the expected stimulus (Rohenkohl & Nobre, 2011; Van Diepen, Cohen, Denys, & Mazaheri, 2015). On the other hand, it has been shown that the temporal expectation of a stimulus improved the perceptual performance by tuning the pre-stimulus alpha oscillation into an optimal phase (Busch, Dubois, & Vanrullen, 2009; Samaha, Bauer, Cimaroli, & Postle, 2015). Of note, there is also debate concerning whether alpha amplitude or alpha phase was responsible for the temporal expectation and the enhanced sensorimotor processing (van Diepen et al., 2015). One account to settle the debate while considering the two-fold results is that the amplitude decrease and the phase concentration of pre-stimulus alpha activity play dissociable roles in different forms of temporal expectation. To assess this account, we investigated how the amplitude decrease and the phase shift of alpha oscillation would be affected by the rhythm-based and sequence-based temporal expectation.

In the present study, we asked how the nested temporal expectation by rhythmic and sequence regularity affected the perceptual performance of the expected stimulus. For this purpose, a stimuli stream with both rhythmic and sequence regularity was created in contrast to an irregular stream. Importantly in the regular stream, a target stimulus was presented in a repeated sequence and meanwhile could be either at an optimal phase (S+P+) or at an antiphase (S+P-) of the entrained neural activity. In a third condition, the target was presented at an optimal phase of the entrained neural activity but was not in a repeated sequence (S-P+). We predicted that relative to S+P- and S-P+ conditions, the perceptual performance would be improved in the S+P+ condition, which benefited from both the rhythmic regularity and the sequence regularity. A perceptual performance engages multiple cognitive modules. In a common perceptual decision-making task, the perceptual attribute has to be recognized and transformed into motor responses based on the stimulus-response mapping defined by the current task. These cognitive modules can be estimated with computational modelling such as the drift-diffusion model (DDM) (Ratcliff, Smith, Brown, & McKoon, 2016), and were suggested to be supported by the neural oscillation during the temporal expectation (Samaha, Iemi, Haegens, & Busch, 2020). With the combination of the DDM, here we elucidated on which cognitive module(s) the nested temporal expectation of rhythmic and sequence regularity can act to improve the perceptual performance. At the neural level, electroencephalographical (EEG) activity was simultaneously recorded to uncover the neural mechanism. Crucially, we investigated how the low-frequency neural entrainment communicated with the pre-stimulus alpha activity, and how amplitude decrease and phase shift of pre-stimulus alpha activity were distinctly affected by the nested structure of temporal expectation.

## Results

Two types of visual stimuli were presented: a regular stream and an irregular stream. In each regular unit, stimuli (each was a disk of Gaussian noise surrounded by a blue ring) were presented in two alternating frequencies (i.e., 1.25 Hz, 2.5 Hz), where five 1.25 Hz-stimuli were followed by five 2.5 Hz-stimuli (Figure 1A). In 75% of the units, a near-threshold target (a Gabor patch surrounded by a red ring) was presented, and participants were required to make a discriminative response to the orientation of the target. For the purpose of our research question, three critical temporal positions were included as the target position: S+P-, S+P+, and S-P+. For S+P-, the first stimulus of the 2.5Hz sequence was presented as the target, by which the target was highly predictable because of the fixed sequence structure while nevertheless being located at an antiphase of the neural oscillation entrained by the preceding 1.25Hz sequence. For S+P+, the last stimulus of the 2.5Hz sequence was presented as the target, by which the target was also highly predictable while being located at an optimal phase of the neural oscillation entrained by the 2.5 Hz sequence. For S-P+, an extra stimulus after the 2.5 Hz sequence was presented as the target, by which the target was unpredictable while being located at an optimal phase of the neural oscillation entrained by the 2.5 Hz sequence. In the irregular stream, stimuli were presented with varying inter-stimulus intervals, but the number of the stimuli preceding the target was matched with each condition in the regular stream.

**Figure 1.**
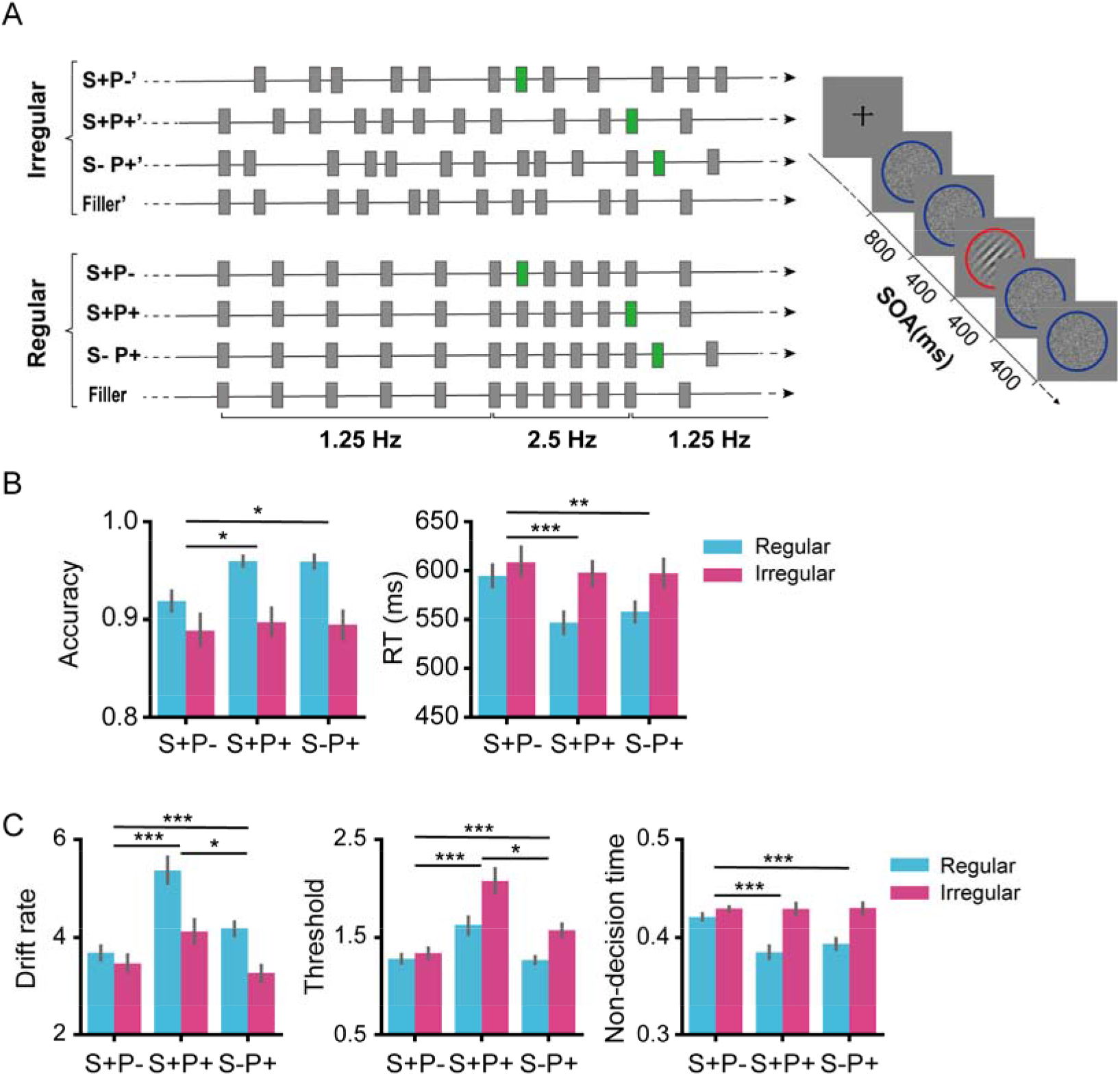
Experimental design and behavioral results. (**A)** Left: the example of stimuli stream in different experimental conditions. The gray bars indicate non-target stimuli and the green bards indicate target stimuli. Right: the sequence of stimuli in an example trial. The target was a near-threshold Gabor patch surrounded by a red ring, and the non-target stimuli were noise surrounded by a blue ring. Participants were asked to make a discriminative response to the orientation of the target. (**B**) Accuracies (left) and reaction times (RTs, right) are shown as a function of experimental conditions. (**C)** Parameters estimated in the HDDM are shown as a function of experimental conditions. Error bars indicate SEM across participants. \**p* < 0.05, \*\**p* < 0.01, *\*\*\*p* < 0.001.

### Accuracy and RTs

A 2 (regularity: regular vs. irregular) × 3 (target position: S+P-, S+P+, vs. S-P+) repeated-measures ANOVA revealed that the accuracy was higher in the regular condition (94.6%) than in the irregular condition (89.4%), *F*(1, 23) = 25.74, *p* < 0.001, *η_p_^2^* = 0.528 (Figure 1B, left). The main effect of position was significant, *F*(2, 46) = 8.58, *p* < 0.001, *η_p_^2^* = 0.272. The accuracy was lower at S+P- (90.4%) than the accuracies at S+P+ (92.9%), *p_bonferroni_* = 0.007, and S-P+ (92.7%), *p_bonferroni_* = 0.023, whereas the difference between S+P+ and S-P+ did not reach significance, *p_bonferroni_* > 0.999. There was an interaction between regularity and target position, *F*(2, 46) = 5.55, *p* = 0.007, *η_p_^2^* = 0.194. Further tests showed that the increased accuracy by regularity was larger at S+P+ (accuracy difference between regular and irregular conditions: 6.3%) than that at S+P- (3.0%), *t*(23) = 3.17, *p_bonferroni_* = 0.013. The increased accuracy by regularity was larger at S-P+ (6.5%) than that at S+P-, *t*(23) = 2.78, *p_bonferroni_* = 0.032. However, the difference between S+P+ and S-P+ did not reach significance, *t* < 1.

The ANOVA on RTs showed that responses were faster in the regular condition (566ms) than the responses in the irregular condition (601ms), *F*(1, 23) =15.9, *p* = 0.001, *η_p_^2^* = 0.408, (Figure 1B, right). The main effect of position was also significant, *F*(2, 46) = 34.5, *p* < 0.001, *η_p_^2^* = 0.600. Pair-wise comparisons showed that responses were faster at S+P+ (572ms), *p_bonferroni_* < 0.001, and S-P+ (578ms), *p_bonferroni_* < 0.001, than responses at S+P- (602ms), whereas the difference between S+P+ and S-P+ did not reach significance, *p_bonferroni_* = 0.141. There was a significant interaction between regularity and position, *F*(2, 26) = 19.3 *p* < 0.001, *η_p_^2^* = 0.456. Further tests showed that the facilitated response by regularity was larger at S+P+ (RT difference between irregular and regular conditions: 51ms) than that at S+P- (14ms), *t*(23) = 6.31, *p_bonferroni_* < 0.001, and larger at S-P+ (39ms) than S+P-, *t*(23) = 3.60, *p_bonferroni_* = 0.005. However, the difference between S+P+ and S-P+ did not reach significance, *t*(23) = 2.24, *p_bonferroni_* = 0.105.

### Hierarchical drift-diffusion modeling (HDDM)

We further modelled the behavioral data with the HDDM (Wiecki, Sofer, & Frank, 2013) to elucidate how the multiple cognitive components were affected by the rhythmic and sequence regularity. The HDDM estimated three parameters: drift rate, decision threshold, and non-decision time. Specifically, the drift rate quantified the accumulation speed of the sensory evidence; the decision threshold quantified the criteria for the perceptual decision; the non-decision time quantified the combined time of early stimulus encoding and the late motor implementation of the decision (Wiecki, Sofer, & Frank, 2013). Convergence was achieved for all of the estimated DDM parameters, R-hats < 1.01.

The 2 × 3 ANOVA revealed that the drift rate was higher in the regular condition (4.41) than in the irregular condition (3.62), *F*(1, 23) = 21.5, *p* < 0.001, *η_p_^2^* = 0.483 (Figure 1C, left). The main effect of position was significant, *F(2,* 46) = 18.9, *p* < 0.001, *η_p_^2^*= 0.451. Pair-wise comparisons showed that the drift rate was higher at S+P+ (4.75) than the drift rates at S+P- (3.57), *p_bonferroni_* < 0.001, and S-P+ (3.72), *p_bonferroni_* < 0.001, whereas no significant difference between S+P- and S-P+ was observed, *p_bonferroni_* = 0.630. The interaction between regularity and position was significant, *F*(2, 46) = 30.6, *p* < 0.001, *η_p_^2^*= 0.571. Further tests showed that the enhanced drift rate by regularity was higher at S+P+ (1.25) than S+P- (0.22), *t*(23) = 6.82, *p_bonferroni_* < 0.001, and S-P+ (0.92), *t*(23) = 2.85, *p_bonferroni_* = 0.027, and the enhancement was also stronger at S-P+ than S+P-, *t*(23) = 5.24, *p_bonferroni_* < 0.001.

The ANOVA showed that the decision threshold was higher in the irregular condition (1.66) than in the regular condition (1.39), *F*(1, 23) = 17.1, *p* < 0.001, *η_p_^2^=* 0.426 (Figure 1C, middle). The main effect of position was significant, *F*(2, 46) = 15.1, *p* < 0.001, *η_p_^2^=* 0.396. Pair-wise comparisons showed that the decision threshold was higher at S+P+ (1.85) than the thresholds at S-P+ (1.42), *p_bonferroni_* = 0.002, and S+P- (1.31), *p_bonferroni_* = 0.001, whereas no significant difference between S+P- and S-P+ was observed, *p_bonferroni_* > 0.167. The interaction between regularity and position was significant, *F*(2, 46) = 22.8, *p* < 0.001, *η_p_^2^* = 0.498. Further tests showed that the lowered threshold by regularity was larger at S+P+ (difference of threshold between irregular and regular conditions: 0.45) than that at S+P- (0.06), *t*(23) = 5.59, *p_bonferroni_* < 0.001, and S-P+ (0.30), *t*(23) = 2.67, *p_bonferroni_* = 0.041, and the threshold decrease was larger at S-P+ than S+P-, *t*(23) = 5.07, *p_bonferroni_* < 0.001.

The ANOVA on non-decision time showed that the non-decision time was shorter in the regular condition (0.400) than in the irregular condition (0.429), *F*(1, 23) = 28.2, *p* < 0.001, *η_p_^2^*= 0.551, (Figure 1C, right). The main effect of position was significant, *F*(2, 46) = 7.0, *p* = 0.002, *η_p_^2^*= 0.233. Pair-wise comparisons showed that the non-decision time was longer at position S+P- (0.425) than that at position S+P+ (0.407), *p_bonferroni_* < 0.014, whereas the difference between position S+P- and S-P+ (0.412), *p_bonferroni_* = 0.074, and the difference between S+P+ and S-P+, *p_bonferroni_* = 0.533, did not reach significance. The interaction between regularity and position was significant, *F*(2, 46) = 25.6, *p* < 0.001, *η_p_^2^*= 0.527. Further tests showed that the facilitated non-decision time by regularity was larger at S+P+ (difference of non-decision time between irregular and regular conditions: 0.044), *t*(23) = 6.51, *p_bonferroni_* < 0.001, and S-P+ (0.036), *t*(23) = 5.98, *p_bonferroni_* < 0.001, than the facilitated non-decision time at S+P- (0.009), whereas the difference between S+P+ and S-P+ did not reach significance, *t*(23) = 1.41, *p_bonferroni_* = 0.516.

### The low-frequency neural oscillation entrained by the rhythmic stimuli

We have assumed that the rhythmic stimuli would induce low-frequency (i.e., 1.25Hz for S+P-, 2.5Hz for S+P+ and S-P+) neural entrainment over the visual cortex. To check this assumption, we calculated the phase-locking value (PLV) for each target position. As shown in Figure 2A, the 1.25 Hz PLV at S+P-, the 2.5 Hz PLV at S+P+ and S-P+ were higher in the regular condition than in the irregular condition (*paired-T* test, *p* < 0.05, with *cluster-based permutation correction*), demonstrating the typical neural entrainment synchronized with the rhythmic stimuli. A data-driven spectral analysis also confirmed the neural entrainment by showing higher amplitudes of the entrained frequencies in the regular condition than in the irregular condition (Supplementary Figure S1). Moreover, the 1.25 Hz phase of S+P- was at an opposite phase (antiphase) of an optimal phase predicted by the 1.25 Hz neural entrainment, as shown by the phase difference (centered mean = 2.94) between S+P- and its preceding position, *p* < 0.001(*Rayleigh Test,* radian, Figure 2B left). By contrast, the 2.5 Hz phases of S+P+ and S-P+ were both at an optimal phase predicted by the 2.5 Hz neural entrainment. The phases were the same, as shown by the phase difference (centered mean = - 0.04) between these two positions, *p* < 0.001(*Rayleigh Test,* radian, Figure 2B right).

**Figure 2.**
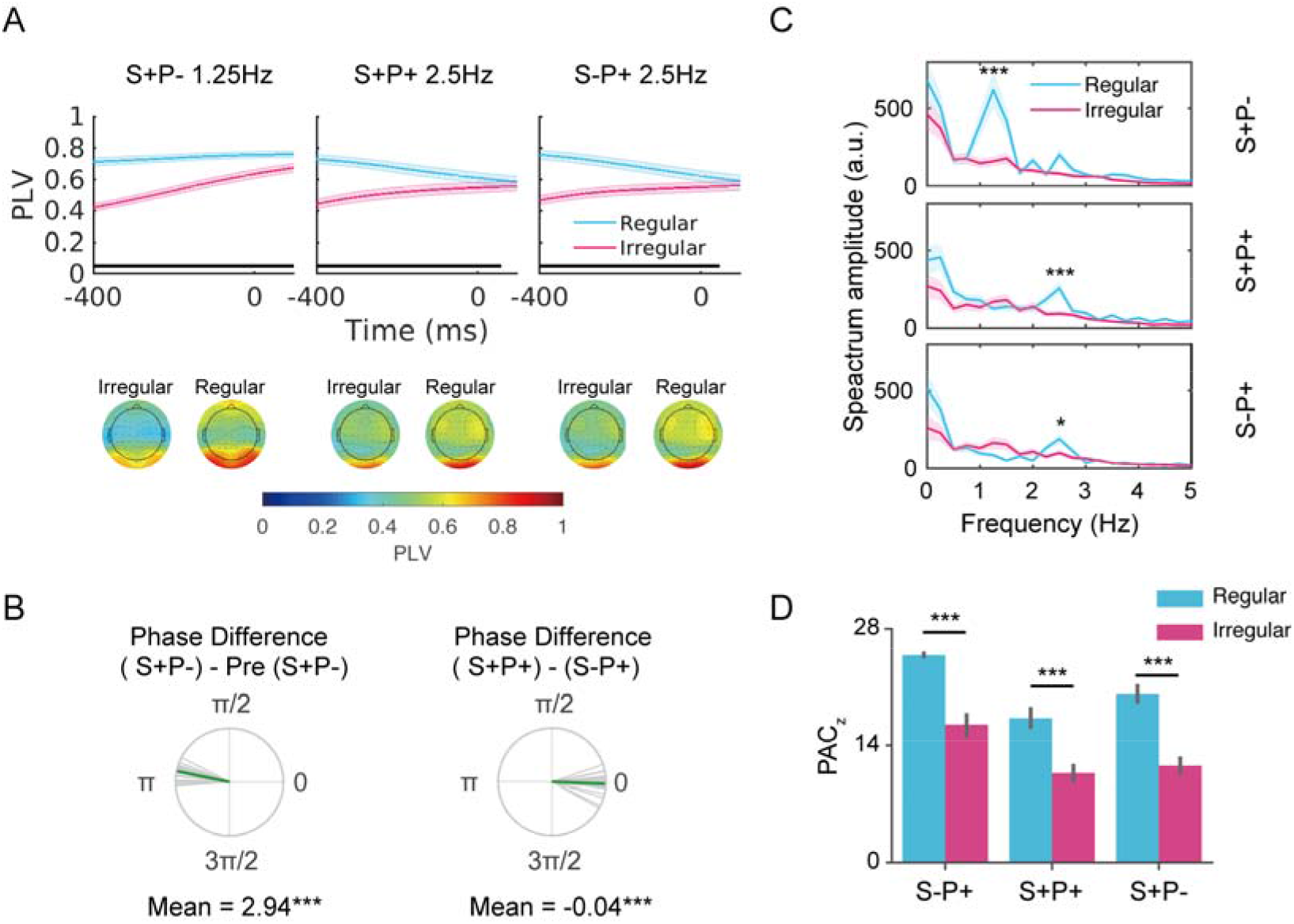
(**A)** Upper panel: the phase locking values (PLV) are shown as a function of the time relative to the target onset in each experimental condition. The black line at the bottom of each graph indicates the time range where the PLV showed a significant difference between regular and irregular conditions, *cluster-based permutation corrected p < 0.05.* Bottom panel: the topographical distribution of the PLV at target onset. (**B)** Left panel: 1.25Hz phase difference between S+P- and the previous position of S+P- (Pre S+P-). The phase differences were clustered around 2.94 (radian), *Rayleigh test, ***p < 0.001.* Right panel: 2.5Hz phase difference between S+P+ and S-P+. The phase differences were clustered around −0.04 (radian), *Rayleigh test, ***p < 0.001.* The green bar indicates the mean phase difference across participants. Each gray bar indicates the phase difference for a specific participant. (**C)** Spectrum amplitudes of alpha amplitude are shown as a function of the low frequencies (0-5Hz) for each experimental condition. At the entrained low frequency, alpha amplitudes showed significantly higher periodicity in the regular condition than in the irregular condition. S+P-: 1.25 Hz; S+P+ and S+P-: 2.5 Hz. The shadows indicate SEM across participants. \**p* < 0.05, \*\*\**p*< 0.001. (**D)** Phase-amplitude coupling (PAC) between the phase of entrained frequency and alpha amplitude for each condition. Error bars indicate SEM across participants.

### Communication between low-frequency entrainment and alpha activity

The fluctuation of the alpha activity (8-12Hz) before the target onset showed a typical periodical characteristic in the regular condition relative to the irregular condition for all of the three positions (Supplementary Figure S2). The FFT analysis showed that the strongest fluctuation of the alpha amplitudes was at 1.25Hz for S+P-, regular vs. irregular: *t*(23) = 5.16, *p*_FDR_ < 0.001, at 2.5 Hz for S+P+, *t*(23) = 6.04, *p*_FDR_ < 0.001, and at 2.5 Hz for S-P+, *t*(23) = 3.84, *p*_FDR_ = 0.012 (Figure 2C). These results suggested that the alpha amplitude was modulated by the phase of the low-frequency neural oscillation. To verify this cross-frequency coupling, we further calculated phase-amplitude coherence (PAC) between the phase of the low-frequency oscillation and the amplitude of the alpha activity. The PACz (Z scored PAC value) was stronger in the regular condition than in the irregular condition for all of the three positions: *t*(23) = 6.54, *p* < 0.001 at S+P- (1.25 Hz), *t*(23) =5.02, *p* < 0.001 at S+P+ (2.5Hz), and *t*(23) = 6.90, *p* < 0.001 at S-P+ (2.5 Hz) (Figure 2D).

### Pre-target alpha amplitude under different structures of temporal expectation

For the analysis of alpha amplitude, the evoked component was firstly removed from the data to avoid the confounding effect contributed by phase (see Materials and Methods). As shown in Figure 3A, the amplitude of the pre-stimulus alpha activity over the visual cortex was higher in the regular condition than in the irregular condition at S+P-, with a significant temporal cluster of −400 to −290 ms relative target onset (*cluster-based permutation corrected p* < 0.001). By contrast, the alpha amplitude was lower in the regular condition than in the irregular condition at position S+P+ and S-P+ (both *p* < 0.001 with *cluster-based permutation correction):* a significant temporal cluster of −400 to 480ms at S+P+, and a significant temporal cluster of −400 to 512ms at S-P+. The reversed pattern of alpha amplitude between S+P- and S+P+/S-P+ conditions suggested that the alpha amplitude was mainly modulated by the phase of the low-frequency oscillation, but was not additionally modulated by the sequence-based expectation. To further test this hypothesis, we compared the amplitude difference between regular and irregular conditions at S+P- with the amplitude difference between irregular and regular conditions (i.e., the reverse of the difference between regular and irregular conditions) at S+P+/S-P+. For each position, the alpha amplitudes were extracted from the significant cluster to calculate the mean difference between regular and irregular conditions (Figure 3B). To avoid making statistical inferences based on non-significant*p* values, we performed the Bayes Factor analysis to quantify the likelihood of the null hypothesis against the alternative hypothesis (Keysers, Gazzola, & Wagenmakers, 2020). The results showed that the hypothesis “the *deceased* amplitude at S+P+ was equivalent to the *decreased* amplitude at S-P+” was 4.46 times more likely to be true than the alternative hypothesis “the decreased amplitude at S+P+ was different from the decreased amplitude at S-P+”. Moreover, the hypothesis “the *decreased* amplitude at S+P+ was equivalent to the *increased* amplitude at S+P-” was 3.60 times more likely to be true than the alternative hypothesis “the decreased amplitude at S+P+ was different from the increased amplitude at S+P-”.

**Figure 3.**
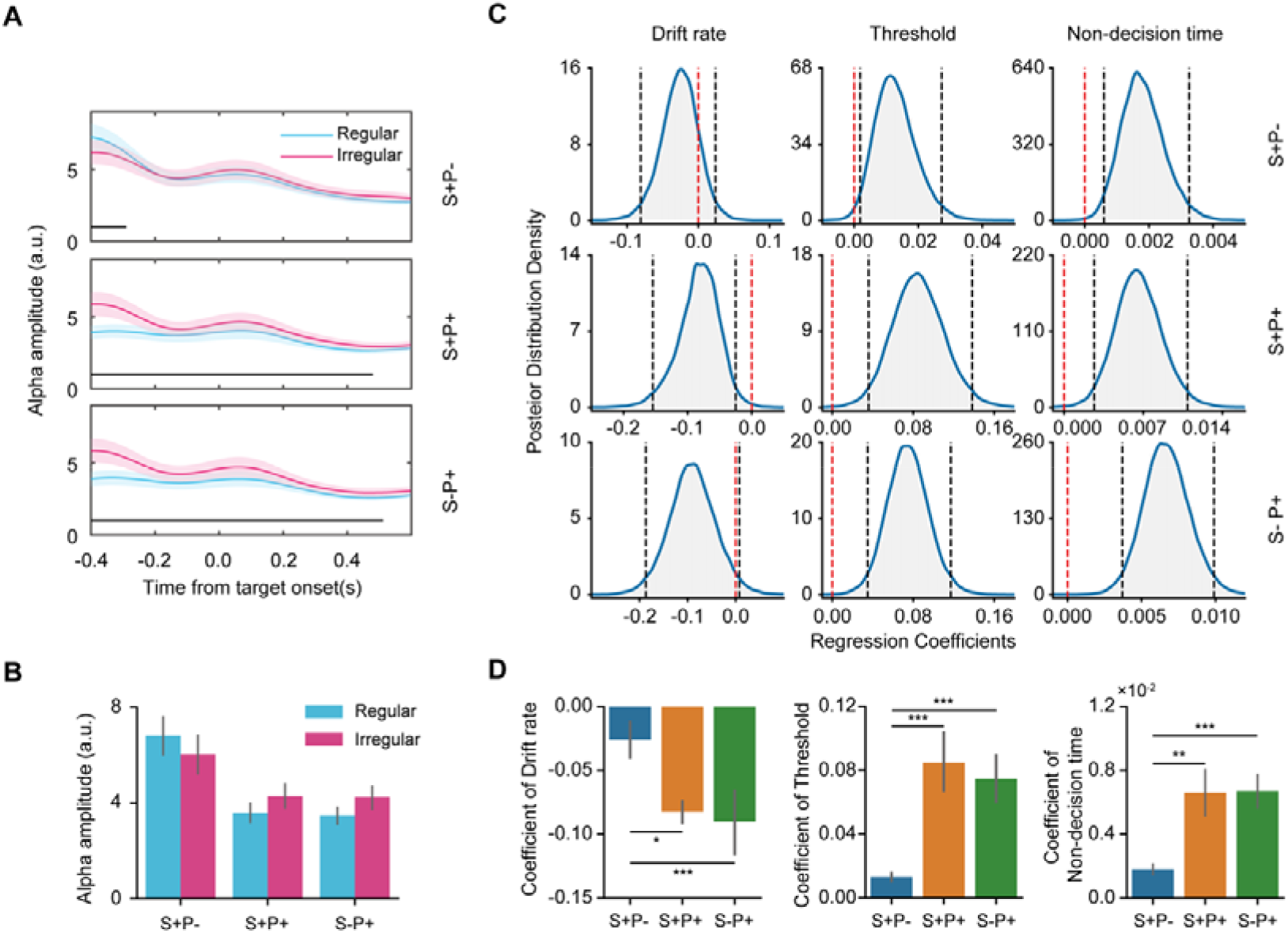
Results of alpha amplitudes. (**A)** Alpha amplitudes are shown as a function of time points relative to the target onset for each experimental condition. The black bar at the bottom of each graph indicates a significant difference between regular and irregular conditions (*cluster-based permutation correctedp < 0.05).* The shadows denote SEM across participants. (**B)** Alpha amplitudes averaged over the significant cluster in (**A)** are shown as a function of the experimental conditions Error bars indicate SEM across participants. To avoid ‘double-dipping’, we did not test the difference between regular and irregular conditions, but rather compared the absolute difference (regular vs. irregular) of amplitudes between positions (see Results). (**C)** The group-level posterior distribution of the regression coefficient was estimated with the regression model for each target position and each one of the HDDM parameters. The regression model quantified to which extent the HDDM parameters could be predicted by alpha amplitude. The black dashed line denotes a 95% confidence interval, and the red dashed line denotes 0 point. (**D)** The individual-level regression coefficients (mean value with SEM) estimated with the regression model are shown as a function of the target position. Error bars indicate SEM across participants. *\*p* < 0.05, *\*\*p* < 0.01, *\*\*\*p* < 0.001.

To assess if and how the amplitude of pre-target alpha activity could be related to the perceptual performance of the target, we tested if the HDDM parameters (drift rate, threshold, and non-decision time) could be predicted by the alpha amplitude. For each position and each HDDM parameter, a regression model that combined with the HDDM was constructed to estimate to which extent the amplitude difference between regular and irregular conditions could predict the difference of the HDDM parameter between regular and irregular conditions. Convergence was achieved for all estimated regression coefficients, R-hats < 1.01. The results of the regression models are shown in Figure 3C. The drift rate could be negatively predicted by the alpha amplitude at S+P+ (mean regression coefficient = −0.082), *p* (*coefficient* > 0) = 0.004, with the drift rate increasing linearly as the alpha amplitude was decreased by regularity (Figure 3C, left). The predictability did not reach significance at S+P- (mean regression coefficient = −0.026), *p* (*coefficient* > 0) = 0.152, whereas was marginally significant at S-P+ (mean regression coefficient = −0.09), *p (coefficient > 0)* = 0.034 (type-I error threshold of 0.025 given the two directions of correlation, see Materials and Methods). In addition, the ANOVA on the coefficients showed a main effect of position, *F*(2, 46) = 7.18, *p* = 0.002, which was due to higher predictability at S+P+ (*p_bonferroni_* < 0.001) and S-P+ (*p_bonferroni_* = 0.012) than at S+P-. No significant difference was observed between S+P+ and S-P+, *p_bonferroni_* > 0.999 (see Figure 3D, left).

The decision threshold could be positively predicted by the alpha amplitude at all of the three positions: S+P- (mean regression coefficient = 0.013), *p (coefficient > 0)* = 0.990), S+P+ (mean regression coefficient = 0.085), *p (coefficient > 0)* > 0.999), and S-P+ (mean regression coefficient = 0.075), *p (coefficient > 0)* > 0.999 (see Figure 3C, middle), with the decision threshold decreasing linearly as the alpha amplitude was decreased by regularity. In addition, the ANOVA on coefficients showed a main effect of position, *F*(2, 46) = 15.0, *p* < 0.001. The predictability was higher at S+P+, *p_bonferroni_* = 0.001 and S-P+, *p_bonferroni_* < 0.001 than S+P-. No significant difference was observed between S+P+ and S-P+, *p_bonferroni_* > 0.999 (Figure 3D, middle).

The non-decision time could be positively predicted by the alpha amplitude at all of the three positions: S+P- (mean regression coefficient = 0.002), *p (coefficient > 0)* = 0.999, S+P+ (mean regression coefficient = 0.007), *p (coefficient > 0)* = 0.999, and S-P+ (mean regression coefficient = 0.007, *p (coefficient > 0)* > 0.999 (Figure 3C, right), with the non-decision time decreasing linearly as the alpha amplitude was decreased by regularity. In addition, the ANOVA on the coefficients showed a main effect of position, *F*(2, 46) = 13.6, *p* < 0.001. The predictability was higher at S+P+, *p_bonferroni_* = 0.004 and position S-P+, *p_bonferroni_* < 0.001 than S+P-. No significant difference was observed between S+P+ and S-P+, *p_bonferroni_* > 0.999 (Figure 3D, right).

The results of the alpha amplitude and the modelling suggested that the alpha amplitude and its contribution to the perceptual performance were mainly driven by the rhythm-based expectation. Even at S+P+ when both the rhythmic and sequence information availed to form temporal expectation, the alpha amplitude was not additionally modulated by the sequence information on top of the rhythmic information. At S+P-, due to the antiphase of the low-frequency oscillation, the alpha amplitude was increased by the rhythmic regularity. The increased alpha amplitude by rhythmic regularity could have been linked to an impaired perceptual performance, which contradicted the observation of improved perceptual performance. This raised the involvement of the sequence-based expectation in improving the perceptual performance, which could be supported by the phase effect of the pre-target alpha oscillation.

### Pre-target alpha phase under different structures of temporal expectation

First, we tested if the phase of the pre-target alpha oscillation was affected by the sequence-based expectation. Considering that the rhythm-based and the sequence-based expectation may have entangling effects, we compared the alpha phase during the time interval before the target (pre-target alpha phase, regular condition) with the alpha phase prior to the stimulus immediately before the target (pre-pre-target alpha phase, regular condition) for each of the three positions. During these two intervals, the neural entrainment induced by the rhythmic stimuli was locked to the same low-frequency phase so that any observed difference should be attributed to the sequence-based expectation. To consider the different lengths of the two intervals at S+P-, the comparison was performed on the 400ms range that was time-locked to the onset of the stimulus before the two intervals (i.e., the onset of the pre-target stimulus for the pre-target alpha phase, the onset of the pre-pre-target stimulus for pre-pre-target phase). At S+P-, the alpha phases showed significant differences during the time interval of 324 to 400 ms relative to the stimulus onset, *p* < 0.001 (Watson–Williams test, with cluster-based correction) (Figure 4A). At S+P+, the alpha phases showed significant differences during the time interval of 320 to 382 ms relative to the stimulus onset, *p* < 0.001 (with cluster-based correction). At S+P+, however, no significant difference in the alpha phase was observed. These results suggested that sequence-based expectation (S+P-, S+P+) of the target changed the pre-target alpha phase.

**Figure 4.**
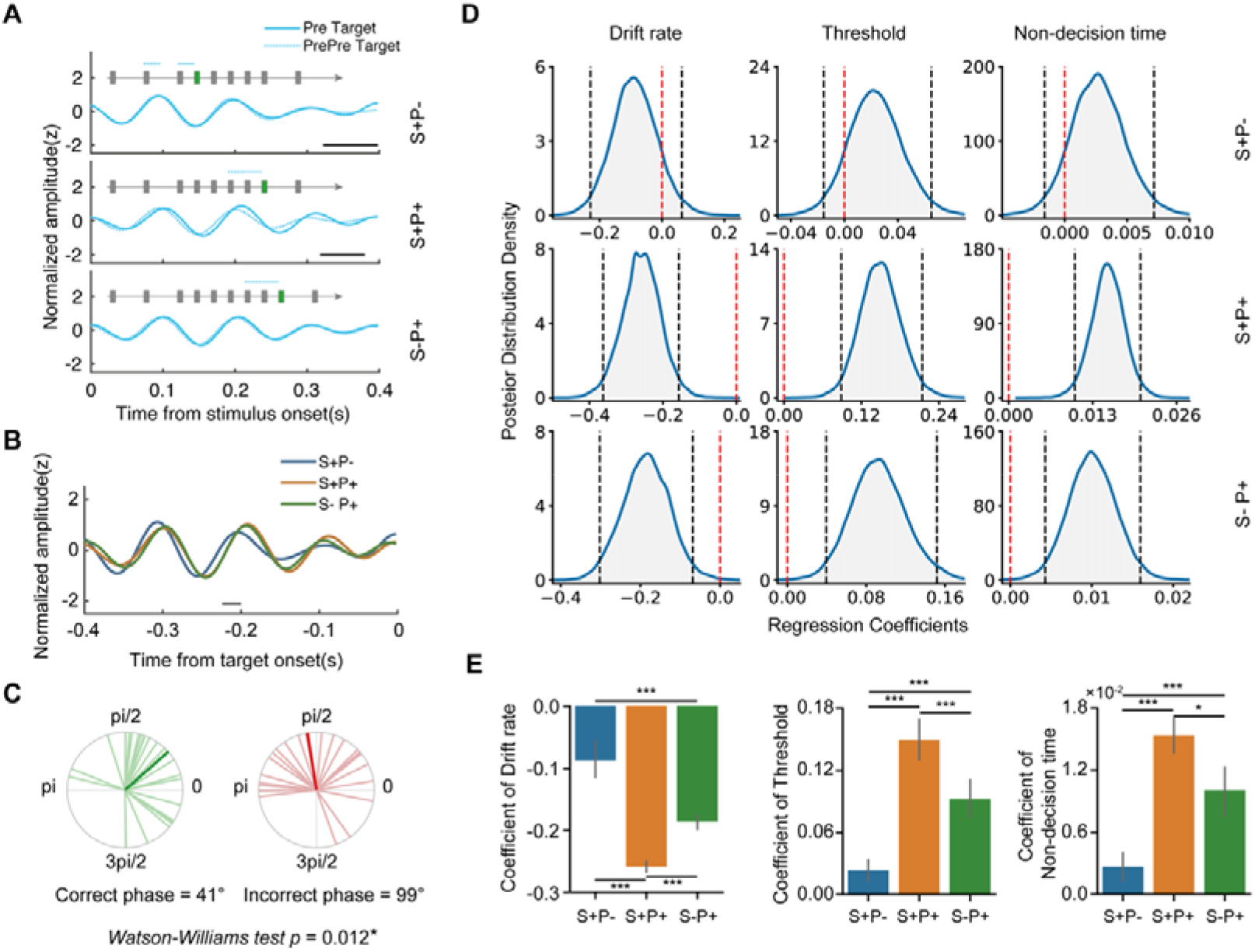
Results of the alpha phase. (**A)** Filtered and normalized alpha amplitudes in the regular condition are shown as a function of time. For each target position (S+P-, S+P+, and S-P+), two time-courses are shown and were compared: the time-course locked to the onset of the stimulus immediately before the target (Pre-Target), and the time course locked to the onset of the stimulus two positions before the target (PrePre-Target). The example of the stimuli stream on the top of each graph illustrates the temporal locations of the two time-courses. The black bar at the bottom of the graph indicates the temporal cluster with a significant difference between the two time-courses, *cluster-based permutation* corrected. (**B)** Filtered and normalized alpha amplitudes in the regular condition are shown as a function of time and target position. Each time course was locked to the target onset. The black bar indicates the temporal cluster with a significant difference between S+P- and S+P+. (**C)**, Alpha phases of trials (collapsed over all experimental conditions) with correct responses (in green color) and trials with incorrect responses (in red color). Each thin bar with light color indicates the alpha phase for a specific participant, and the thick bar with dark color indicates the mean of the alpha phase across participants (**D)** The group-level posterior distribution of the regression coefficient estimated with the regression model for each target position and each one of the HDDM parameters. The regression model quantified to which extent the HDDM parameters could be predicted by the distance between the alpha phase at the target onset and the optimal alpha phase. The black dashed line denotes a 95% confidence interval, and the red dashed line denotes 0 point. (**E)** The individual-level regression coefficients (mean value with SEM) estimated with the regression model are shown as a function of the target position. Error bars indicate SEM across participants. *\*p* < 0.05, *\*\*p* < 0.01, *\*\*\*p* < 0.001.

Then, we tested if the rhythm-based expectation also contributed to the pre-target alpha phase by comparing the phases during the pre-target interval between different conditions. The results showed significant differences in the alpha phase between S+P- and S+P+ during the time interval of −224 to −200 ms relative to the target onset, *p* < 0.001 (Watson–Williams test, with cluster-based correction) (Figure 4B). However, no significant difference was observed between S+P- and S-P+, or between S+P+ and S-P+. The difference between S+P- and S+P+ suggested that the pre-target alpha phase was also affected by rhythm-based expectation.

Last but not least, we assessed if and how the alpha phase was biased by the temporal expectation to improve the perceptual performance. We first sorted the data into epochs with a correct response and epochs with an incorrect response, with the expectation that the target onset of the former would be more likely to be localized at an optimal phase than the latter (Samaha et al., 2015). Consistent with this assumption, the epochs with correct responses and the epochs with incorrect responses showed a significant difference in the alpha phase, *Watson-Williams test p =* 0.012 (Figure 4C). We thus defined the mean phase of the correct responses as the optimal phase and calculated the phase distance in terms of the circular distance between the phase at the target onset in each trial and the optimal phase. To avoid the risk of double-dipping, we did not test the difference in phase distance. Instead, we focused on the predictability of the phase distance on the single-trial perceptual performance.

Similar regression models to the above were conducted to assess to which extent the HDDM parameters could be predicted by the alpha phase. Convergence was achieved for all regression coefficients, R-hats < 1.02. The drift rate could be negatively predicted by the phase distance at S+P+ (mean regression coefficient = −0.259), *p (coefficient > 0)* < 0.001, and S-P+ (mean regression coefficient = −0.186), *p (coefficient > 0)* = 0.002, (Figure 4D, left), with the drift rate increasing linearly as the alpha phase was closer to the optimal alpha phase, whereas the predictability at S+P- (mean regression coefficient = −0.087) did not reach significance, *p (coefficient > 0)* = 0.114. The ANOVA on the coefficients showed a main effect of position, *F*(2, 46) = 30.8, *p* < 0.001, which was due to higher predictability at S+P+ (*p_bonferroni_* < 0.001) and S-P+ (*p_bonferroni_* < 0.001) than S+P-, and also higher predictability at S+P+ than S-P+, *p_bonferroni_* < 0.001 (Figure 4E, left).

The decision threshold could be positively predicted by the phase distance at S+P+ (mean regression coefficient = 0.149), *p (coefficient > 0)* > 0.999, and S-P+, (mean regression coefficient = 0.093), *p (coefficient > 0)* > 0.999, (see Figure 4D, middle), with the decision threshold decreasing linearly as the alpha phase was closer to the optimal phase, but not at S+P- (mean regression coefficient = 0.023), *p (coefficient > 0)* = 0.882. The ANOVA on the coefficients showed a main effect of position, *F*(2, 46) = 42.1, *p* < 0.001. The predictability was higher at S+P+ than S+P- (*p_bonferroni_* < 0.001) and S-P+ (*p_bonferroni_* < 0.001), and was also higher at S-P+ than S+P-, *p_bonferroni_* < 0.001 (Figure 4E, middle).

The non-decision time could be positively predicted by the phase distance at position S+P+ (mean regression coefficient = 0.015), *p (coefficient > 0)* > 0.999, and S-P+ (mean regression coefficient = 0.010), *p (coefficient > 0)* > 0.999 (see Figure 4D, right), with the non-decision time decreasing linearly as the alpha phase was closer to the optimal phase, but not at S+P- (mean regression coefficient = 0.003), *p (coefficient > 0)* = 0.895. In addition, the ANOVA on the coefficients showed a main effect of position, *F*(2, 46) = 31.0, *p* < 0.001. The predictability was higher at S+P+ than S+P-, *p_bonferroni_* < 0.001) and S-P+, *p_bonferroni_* = 0.014, and was also higher at S-P+ than S+P-, *p_bonferroni_* = 0.001 (Figure 4E, right).

## Discussion

In the present study, we investigated how rhythmic and sequence information can be combined to form temporal expectation and optimize the perceptual performance of the expected stimulus. The behavioral results showed that the rhythm-based and sequence-based expectation had an additive effect in improving the perceptual performance of the expected stimulus. The EEG results suggested a dissociation of alpha amplitude and phase in supporting the perceptual performance under different structures of temporal expectation. While the amplitude of pre-stimulus alpha oscillation was mainly driven by the rhythmic information through phase-amplitude coupling, the phase of pre-stimulus alpha oscillation was affected by both the rhythmic and sequence information. At the single-trial level, on the one hand, the rhythm-based expectation improved the perceptual performance by reducing the pre-stimulus alpha amplitude, whereas the sequence-based did not further affect the alpha amplitude and the corresponding perceptual performance. On the other hand, the rhythm-based and the sequence-based expectation had a synergistic effect in biasing the alpha phase toward the optimal phase to improve the perceptual performance.

Mounting evidence has demonstrated that perceptual performance can be improved by different sources of temporal expectation (see Nobre & van Ede, 2018 for a review). For instance, relative to arrhythmic information, the perceptual sensitivity (Cravo et al., 2013) and the motor implementation of the perceptual decision (Rohenkohl & Nobre, 2011) were enhanced by rhythmic information of the stimuli. Relative to an unpredictable stimuli sequence, the perceptual sensitivity of stimuli in a repeated sequence was also enhanced (Morillon et al., 2016). In previous studies, however, the different sources of temporal expectation were often treated in separate task contexts, with a single form of temporal expectation in a specific context (but see Breska & Deouell, 2014 for temporal expectation induced by concurrent rhythm and a predictive cue). In an extension, our HDDM results showed that the rhythmic and sequence information had an additive effect on the perceptual sensitivity to the expected stimulus by both increasing the accumulation of sensory evidence and alleviating the threshold for the perceptual decision. The non-decision time of the perceptual decision, however, benefited more from the rhythm-based expectation, which might be due to more efficient motor implementation as highlighted by the link between the motor system and the rhythmic processing (Cannon & Patel, 2021; Morillon, Schroeder, & Wyart, 2014). These results suggested that the various temporal information in the environment was utilized to affect perceptual decision-making.

It has been well-documented that brain oscillations on multiple time scales are coordinated to modulate sensorimotor processing (Palva & Palva, 2018; Schroeder & Lakatos, 2009). It has been suggested that the momentary power of high-frequency oscillation is determined by the phase of the low-frequency oscillation through cross-frequency coupling (Lakatos et al., 2008). In accordance with this notion, our results showed a strong dependence of the alpha amplitude on the phase of the low-frequency neural entrainment, with the pre-stimulus alpha oscillation fluctuating in the same frequency of the rhythmic stimuli. A corresponding benefit is that a stimulus aligned with the high-excitability phase gains improved perceptual processing, whereas a cost is that a stimulus aligned with the low-excitability is less likely to be efficiently recognized (Lakatos et al., 2008). This prediction was supported by our results that the pre-stimulus alpha amplitude was reduced when the rhythmic stimulus was at an optimal phase of the low-frequency entrainment (i.e., S+P+ and S-P+), and the single-trial perceptual decision-making was critically predicted by the alpha amplitude. However, when the rhythmic stimulus was at an antiphase (i.e., S+P-), the alpha amplitude was otherwise increased, indicating a lowered preparing state for the stimulus. Our results elucidated the mechanism that the rhythmic regularity in the environment affected the perceptual performance through the cross-frequency coupling between the phase of the low-frequency entrainment and the amplitude of pre-stimulus alpha oscillation.

While both the amplitude reduction (Breska & Deouell, 2017; Rohenkohl & Nobre, 2011; Van Diepen et al., 2015) and the shifted phase (Busch et al., 2009; Samaha et al., 2015) of pre-stimulus alpha oscillation have been suggested as neural mechanisms of temporal expectation, an important finding here is the dissociated functions of alpha amplitude and alpha phase in the different structures of temporal expectation. The current results showed that the alpha amplitude was mainly driven by the rhythmic information, whereas the sequence information did not additively modulate the alpha amplitude. This notion is supported by the following evidence: 1) at an optimal phase of the rhythmic stimuli, the decreased alpha amplitude was equivalent regardless of the presence of the sequence-based expectation (S+P+ vs. S-P+); 2) the decreased alpha amplitude at an optimal phase (S+P+) was equivalent to the increased alpha amplitude at an antiphase (S+P-) of the rhythmic stimuli; 3) the predictive power of the alpha amplitude on the single-trial perceptual performance was not additionally contributed by the sequence-based expectation (S+P+ vs. S-P+). In contrast, the alpha phase was modulated by both rhythmic and sequence information, leading to synergistic effects in biasing the alpha oscillation toward an optimal phase where the perceptual performance can be optimized. Specifically, on top of the rhythmic information, the addition of sequence information induced a change in the phase of the pre-target alpha oscillation (i.e., at both S+P- and S+P+), whereas such phase change was not observed when the sequence-based expectation was absent (i.e., at S-P+). And, the phase of the pre-target alpha oscillation was also affected by whether the target was at an optimal phase of rhythmic stimuli (i.e., S+P- vs. S+P+). At the single-trial level, the perceptual performance was predicted by the extent to which the phase was close to the optimal phase of the alpha oscillation, and the combination of the rhythmic and sequence information rendered the highest predictive power.

Although the alpha amplitude was involved only in the rhythm-based expectation here, it should not be generalized into that alpha amplitude was immune to the sequence-based expectation regardless of task context. Instead, the suggestion based on the current findings is that the alpha amplitude and alpha phase were flexibly coordinated to take effect according to the task. The rhythmic processing has been suggested as a default mode of the brain, the high and low excitability of which can be reset on multiple time scales (Jones, Johnston, & Puente, 2006; Schroeder & Lakatos, 2009). Due to this dominant role of the rhythmic processing and that a high proportion of the stimuli in the regular stream was aligned with the high-excitability phase (i.e., onbeat stimuli), it would be economic to have a fixed relationship between alpha amplitude and the phase of the entrained low-frequency oscillation, with the alpha phase being flexibly regulated to achieve efficient processing. Here the alpha phase was shifted both by the phase-resetting of the rhythmic stimuli, and by the expectation acquired from the repeated sequence. The phase shift regulated by sequence-based expectation can add to improving the processing of the onbeat target (i.e., S+P+), and more importantly, overcome the lowered processing of the offbeat target (i.e., S+P-). Similar adjustment by top-down temporal expectation in compensating for the processing of the offbeat targets was also shown in a recent study (Breska & Deouell, 2016). When there was a high probability of offbeat targets such that the onset of the offbeat target can be predicted, the contingent negative variation (CNV), an event-related potential component of expectation (Walter, Cooper, Aldridge, Mccallum, & Winter, 1964), was adjusted toward the expected time of the offbeat target (Breska & Deouell, 2016). The flexible collaboration of alpha phase and alpha amplitude not only helped to settle the debate concerning the alpha oscillation in temporal expectation (van Diepen et al., 2015) but also suggested that the various temporal information was flexibly utilized through the coordination of distinct neural mechanisms to efficiently achieve the task goal.

A nested structure of regularity is fundamental in daily-life activities such as speech and music (Ding, Melloni, Zhang, Tian, & Poeppel, 2015; Koelsch, Rohrmeier, Torrecuso, & Jentschke, 2013). Our findings shed light on the dynamic interaction of neural oscillations in exploiting a nested structure of temporal regularity. This echoes with previous studies of different cognitive contexts (Palva & Palva, 2018). For instance, Yuan et al. (2021) showed that the multiscale rhythmic information in a stimuli stream was used to alleviate the attentional blink (Raymond, Shapiro, & Arnell, 1992), a cognitive “bottleneck” in visual attention, and this behavioral change was critically predicted by the phase-amplitude coupling between the multiscale neural oscillations that correspond to the temporal structure of the stimuli (Yuan, Hu, Zhang, Wang, & Jiang, 2021). Beyond visual perception, the cross-frequency interaction was also found as crucial for working memory (Axmacher et al., 2010) and speech segmentation (Gross et al., 2013). Our results showed that the dynamic interaction of brain oscillations is not only expressed as the cross-frequency coupling (e.g., the phase-amplitude coupling), but also as the amplitude-phase coordination within a specific frequency range (e.g., the alpha oscillation). Taken together, these findings suggested that the multiscale brain oscillations are flexibly organized to deal with the complex environment and empower adaptive behaviour.

## Materials and Methods

### Participants

A total of 26 right-handed university students participated in this study. One participant was excluded from analysis due to incomplete data caused by technical errors during the experiment, and another participant was excluded from data analysis due to excessive artifacts (50% of total trials) of the EEG signals, resulting in 24 participants (10 females, mean age 20.5 years old). All participants had normal or corrected-to-normal visual acuity and normal color vision and reported no history of psychiatric or neurological disorders. Informed consent was obtained from each participant prior to the experiment. This study was carried out in accordance with the Declaration of Helsinki and was approved by the Ethics Committee of the School of Psychological and Cognitive Sciences, Peking University (2019-12-01).

### Stimuli and Apparatus

Stimuli were created and presented using Psychtoolbox-3 extension for MATLAB (Brainard, 1997). Stimuli were presented on a Display++ monitor (1920*1080 spatial resolution, 120Hz refresh rate) against a gray background (RGB:125, 125, 125). The eye-to-monitor distance was fixed at 70 cm. A chin rest was used to maintain the head position and a constant viewing distance. Responses were collected using a standard keyboard.

Stimuli were presented in a stream flow at the center of the screen, one at a time (Figure1A). Two types of stimuli were used in the experiments: standard stimuli and target stimuli. Standard stimuli were disks of Gaussian noise (radius: 2° of visual angle), each of which was surrounded by a blue ring (RGB: 0, 0, 200). The disks were created by smoothing Gaussian patches with a two-dimensional kernel. The smoothing dimension (0.083° of visual degree) and the root-mean-square contrast of the noise patch were fixed across experiments and participants. Each Target stimulus was a Gabor patch surrounded by a red ring (RGB:200, 0, 0), and was embedded in the standard stimuli. The orientation of the Gabor patch was either 135° or 45° relative to the horizontal axis, and the spatial frequency of the Gabor patches was 2 cycles/degree of visual angle. The above-mentioned parameters of the stimuli were chosen following a previous study (Rohenkohl, Cravo, Wyart, & Nobre, 2012). Prior to the experiment, a psychophysical test was conducted to estimate the contrast threshold with 75% accuracy in discriminating the orientation of the Gabor patches, with an adaptive staircase procedure (Kaernbach, 1991). The contrast that 10^0.1^ times of contrast threshold corresponding to 75% accuracy was used in the formal experiment across all conditions.

### Design and procedure

There were two types of stimuli stream: the regular stream and the irregular stream. In both streams, each stimulus remained on the screen for 50 ms. Within the regular stream, stimuli were presented at two alternating frequencies (i.e., 1.25 Hz, and 2.5 Hz). The stimulus onset asynchrony (SOA) between successive stimuli changed with a fixed rule: five SOAs of 800 ms followed by five SOAs of 400 ms which was then followed by five 800 ms SOAs and so on. Every ten stimuli were grouped into a unit where the long-SOA stimuli were always followed by the short-SOA stimuli. Within the irregular stream, the SOA between each successive two stimuli was randomly chosen from 300 ms, 400 ms, 500 ms, 600 ms, 700 ms, 800 ms, and 900 ms. Within the irregular stream, every ten successive stimuli were also grouped into a unit. In each unit, there was a 75% probability that one target stimulus was embedded in the stream while a 25% probability that no target was presented at all (Target absent, filter unit). No target was presented in the very first unit of each trial. In the regular stream, the target was presented at one of the three temporal positions with equal probability (as shown in Figure 1A): 1) the first position of the short-SOA stimuli following the long-SOA stimuli, which could be predicted by the stimuli sequence but not predicted by the rhythm (i.e., 1.25Hz) that preceded the position (S-P+); 2) the last position of the short-SOA stimuli that followed by the long-SOA stimuli in the next unit, which could be predicted by both the stimuli sequence and the rhythm (i.e., 2.5Hz) that preceded the position (S+P+); 3) an extra position inserted after the last position of the short-SOA stimuli that followed by the long-SOA stimuli, with a short SOA before while a long SOA after this extra position, and this position could be predicted by the rhythm (2.5Hz) but not by the stimuli sequence (S-P+). The three positions within the irregular stream (S’+P’-, S’+P’+, S’-P’+) were matched the positions within the regular stream in the way that the numbers of stimuli between two target stimuli were the same in the two streams, and the SOAs before and after the positions within the irregular stream were consistent with the regular stream.

Each trial started with a black central fixation cross (RGB: 0, 0, 0, 1° of visual angle), which remained on the screen until a space key was pressed by the participant to begin the current trial. The stimuli stream was immediately presented after the key press. Participants were asked to respond to the orientation of the target stimulus (left vs. right) with the left and right index fingers respectively. They were asked to make responses as quickly and accurately as possible. A white central fixation cross (RGB: 255, 255, 255, 1° of visual angle) was presented at the end of each trial and remained on the screen for 10s until the next trial started.

In each trial, there were 158 standard stimuli and 12 target stimuli, and the target was presented at one of the 3 positions with equal probability. Units with different target positions were mixed and presented in a pseudorandom order such that no more than two consecutive units had the same target position. There were 20 trials of the regular stream and 20 trials of irregular stream, resulting in 240 targets for each of the two streams. The two types of trials were presented in two separate blocks, with the order of the two blocks counterbalanced across participants. After every 5 trials, there was a 1-minute break.

Therefore, the experiment had a 2 (Regularity: regular vs. irregular) * 3(Target position: S+P-, S+P+, vs. S-P+) within-subject design.

### Statistical analysis of behavioural data

For each participant, incorrect responses, omissions, and responses with RTs beyond mean RT ± 3 standard deviation (1.80%) in each condition were excluded. The mean RT of the remaining responses for each condition was calculated. The accuracy for each condition was calculated as the proportion of the number of correct responses against the total number of targets in each condition. Accuracy and RTs were subjected to a repeated-measure ANOVA with target position and stream regularity as within-subject factors.

The DDM that characterizes the evolving perceptual decisions in a 2-alternative forced-choice task (Stafford, Pirrone, Croucher, & Krystalli, 2020; Sun & Landy, 2016; Tavares, Perona, & Rangel, 2017) was used to model the behavioral data. Here we estimated three parameters, drift rate, threshold, and non-decision time to simulate the perceptual decision-making processes using the HDDM 0.6.0 toolbox (Wiecki et al., 2013). The HDDM model is a hierarchical Bayesian estimation of drift-diffusion parameters and generates parameter estimates at both individual level and group level. Omission trials were excluded prior to modelling and the probability of outlier was set to 5%. In each experimental condition, for the convergence of the parameters, each of the three parameters was estimated with an independent model. Four Markov Monte-Carlo chains were used to estimate the parameters, with 10000 samples in each chain while the first 2000 samples were discarded as burn-in to achieve convergence. We computed the R-hat (Gelman-Rubin) convergence statistics to ensure the convergence of the models (Gelman & Rubin, 1992). Individual parameters were averaged across the remaining 32000 samples for further analysis. The three parameters were subjected to a repeated-measure ANOVA with target position and stream regularity as the within-subject factors.

### EEG recording and preprocessing

EEG signals were recorded by 64 Ag/AgCl electrodes mounted in an elastic cap (Easy-cap Brain products, Germany) according to an extending 10-20 system. Vertical electrooculograms (EOG) were recorded by an electrode placed below the center of the right eye. The impedance of all electrodes was kept below 5 kΩ. The EEG and EOG signals were amplified by two Brain-Amp amplifiers (Brain Product, Germany), digitalized to a sample of 500Hz, and were online filtered by a band-pass filter of 0.016Hz-100Hz. EEG signals were online referenced to the FCz electrode. Preprocessing (denoising and segment) was conducted with EEGLAB toolbox (Delorme & Makeig, 2004). The offline data were band-pass filtered between 0.5Hz-60Hz and re-referenced to the averaged signal of right and left mastoid electrodes. Independent component analysis was performed to remove eye-movement and other artifact components (Drisdelle, Aubin, & Jolicoeur, 2017). Stimulus-locked epochs were extracted from the interval of −4000 to 1500ms relative to the target onset. The long epoch was selected to avoid potential edge effects for the analysis in the frequency domain. Baseline corrections were applied to the interval of −1000ms to 0ms relative to the target onset. Epochs with amplitude exceeding 100 μV were removed for further analysis.

### Phase-locking value (PLV) of low-frequency neural oscillation

To show the neural entrainment by the rhythmic visual stimuli, the phase-locking value (PLV) of the pre-target activity was calculated over the visual cortex. The PLV was expected to be higher in the regular condition than in the irregular condition. For each of the occipital channels (Oz, O1, O2, POz, PO3, PO4, PO7, PO8), the stimulus-locked data (time range: - 4000 to 1500 ms relative to target onset) was filtered using a two-way least-squares FIR filter (position **S**: 0.75Hz-1.75Hz, center frequency 1.25Hz; S+P+ and S-P+: 2.0Hz-3.0Hz, center frequency 2.5Hz. *eegfilt,* EEGLAB). The phase of the filtered data was obtained with Hilbert transformation. For each participant and each condition, the phase-locking value at each time point was calculated as the length of the mean vector of the phases across the targets of a specific condition. For each of the three positions, the difference of PLV between the regular and the irregular condition was tested with *paired-T* test. This test was performed for each time point of the −400 to 100 ms interval relative to the target onset, with *cluster-based permutation* (number of iterations = 1000, minimum time duration = 20ms, cluster level alpha = 0.05) for correcting multiple comparisons across the time points.

To test if S+P- was at an antiphase of an optimal phase of 1.25Hz entrained neural activity, the difference between the phase of 1.25 Hz at the target onset and the phase of 1.25 Hz at the onset of the preceding stimulus before the target was obtained for each participant. The preceding stimulus was expected to be at an optimal phase of the entrained neural activity at 1.25Hz. Rayleigh test was used to test whether the phase difference was uniformly distributed or centered around 180°. For S+P+ and S-P+, the phase difference of 2.5 Hz at the target onset between the two positions was calculated for each participant. The two positions were expected to be at an optimal phase of the entrained neural activity at 2.5 Hz. Rayleigh test was used to test whether the phase difference of 2.5 Hz between S+P+ and S-P+ was uniformly distributed or centered around 0°.

### The extraction of alpha amplitude and alpha phase

For each of the occipital electrodes, the stimulus-locked data (time range: −4000 to 1500 ms relative to target onset) was filtered (8-12 Hz) using a two-way least-squares FIR filter (*eegfilt*, EEGLAB). The filtered data was then transformed with a Hilbert transformation (*hilbert*, MATLAB). The alpha phases were extracted from the transformation and averaged over the occipital electrodes.

To obtain alpha amplitude, in each condition, the average of the epoched data was firstly subtracted from the data epochs. This was to remove the evoked components in the data set so that the confounding effect of alpha phase and alpha amplitude can be avoided. Then the induced epoch was filtered with a two-way least-squares FIR filter (8-12 Hz, *eegfilt*, EEGLAB) and transformed with Hilbert transformation (*hilbert*, MATLAB). The transformed amplitudes were averaged across the occipital electrodes.

### Fast Fourier transformation (FFT) of alpha amplitude and phase-amplitude coupling (PAC) analysis

To investigate if the alpha activity was affected by the rhythmic stimuli and hence would show fluctuations at the low frequency, FFT analysis was performed on the alpha amplitude with the expectation that alpha oscillation would show stronger fluctuation in the regular condition than in the irregular condition, especially at 1.25 Hz and 2.5 Hz (i.e., the frequency of the rhythmic stimuli). The time courses of alpha amplitude (8-12 Hz) were extracted from the interval of −2000 to 0 ms relative to the target onset and averaged across epochs. The spectrum amplitudes (from 0.25Hz to 250 Hz, in steps of 0.25 Hz) of the averaged alpha amplitude were calculated using FFT. The zeros-padding method was used to improve the frequency resolution. For each of the three positions (S+P-, S+P+, S-P+), the spectrum difference between regular and irregular conditions was compared with *Paired-T* tests (one-tailed). FDR correction was applied for multiple comparisons across 0-5 Hz frequencies.

PAC was performed to verify the cross-talk between the low-frequency neural entrainment and the alpha amplitude. For each condition, the alpha (8-12 Hz) amplitude and phase time-courses of the low-frequency activity (1.25 Hz for S+P-; 2.5 Hz for S+P+ and S-P+) were extracted and averaged across epochs. For each time-point during the interval of −2000 to 0 ms relative to target onset, the alpha amplitudes were transformed into z-scores. A vector of phase-amplitude time course was constructed, with the phase of low-frequency entrainment as angle and the amplitude of alpha-band as length. For each condition, the mean vector was calculated over the selected time intervals (−2000 to 0 ms). The length of the mean vector was defined as the PAC value for the low-frequency phase and alpha amplitude. To avoid the bias that may be caused by the non-uniform distribution of the phase, we normalized the PAC value with nonparametric permutations. Specifically, for each condition, each amplitude time course was shuffled across the time interval and the PAC value based on the shuffled data was calculated. This calculation was repeated 1000 times, rendering a distribution of permutated PAC values. The normalized Z value of the PAC (PAC_z_) was then obtained by subtracting the mean of the permutated PAC values from the un-permutated PAC values and divided by the standard deviation of the permutated PAC values. A *paired-T* test was performed to compare the PACz between regular and irregular conditions for each position.

### Analysis of pre-target alpha amplitude

For each of the three positions, the averaged alpha amplitudes across epochs were compared between regular and irregular conditions using a *paired-T* test. This test was performed for each time point of the −400 to 600 ms interval relative to the target onset with *cluster-based permutation* (number of iterations = 1000, minimum time duration = 20ms, cluster level alpha = 0.05) for correcting multiple comparisons across time points. We chose this time range because it covered both the temporal expectation of the target and the sensorimotor processing of the target.

To examine if the alpha amplitude could be related to the perceptual performance, we tested if the HDDM parameters (Drift rate, Threshold, Non-decision time) could be predicted by the alpha amplitude. For each of the three positions, the amplitude difference between the regular and irregular conditions was included in a regression model that was combined with the HDDM model (*HDDMRegressor,* HDDM). By including the regular condition as the intercept (baseline) and the irregular condition as a fixed regressor, the regression model estimated to which extent the three HDDM parameters (drift rate, threshold, non-decision time) difference could be predicted by the amplitude difference between regular and irregular conditions. The regression coefficient was estimated using a hierarchical Bayesian estimation method and the posterior distribution of the regression coefficient was estimated using the Markov chain Monte Carlo techniques (MCMC, 4 chains, 10000 samples for each chain, the first 2000 samples were discarded). The Gelman-Rubin R-hat statistics were used to assess the convergence of the model parameters. For each of the three positions, statistical testing was conducted by calculating the probability of the regression coefficient in the posterior distribution. The significance criterion (Type-I error threshold) was set as 0.05. A *p* > 0.975 suggested that the difference of an HDDM parameter could be positively predicted by the amplitude difference, and *p* < 0.025 suggested that an HDDM parameter could be negatively predicted by the amplitude difference. In addition, to assess if the predictability of phase distance on parameters was affected by different positions, the regression coefficients at different positions were submitted to a repeated-measures of ANOVA.

### Analysis of pre-target alpha phase

For each of the three positions, we tested if the alpha phase after the stimulus preceding the target (i.e., pre-target alpha phase) was different from the alpha phase after the stimulus that two positions before the target (i.e., pre-pre-target alpha phase). This phase difference was conducted in the regular conditions provided that the time intervals were random in the irregular condition. For the pre-target alpha phase, the 400 ms interval was time-locked to the onset of the pre-target stimulus; for the pre-pre-target alpha phase, the 400 ms interval was time-locked to the pre-pre-target stimulus (i.e., the stimulus that two positions before the target). For each time point, the normalized alpha phases (averaged across epochs) were compared with the Watson–Williams test (Watson and Williams 1956; Stephens 1969), a circular analog of the t-test, which tests if the two samples of angles have different phase distributions. Cluster-based permutation correction was used to solve the multiple comparison problem across time points. The alpha phases during the time interval of −400 to 0 ms relative to the target onset at the three positions (regular condition) were also compared with the Watson–Williams test.

For each participant, “optimal” and “non-optimal” alpha phases were differentiated based on the correctness (correct vs. incorrect) of the behavioural response. For each participant, across all epochs (regardless of condition) with correct responses, the alpha phases at the target onset were averaged as the “optimal” phase; and across all epochs with incorrect responses, the alpha phases at the target onset were averaged as the “non-optimal” phase. Watson-Williams test was used to test the phase difference between correct and incorrect responses. For each epoch, the phase distance was calculated as the difference between the phase at the target onset and the “optimal” phase. The same regression models were fitted to estimate to which extent the HDDM parameters could be predicted by the phase distance.

## Acknowledgments

This study is supported by the National Natural Science Foundation of China (32000779, 31861133012) and the Shanghai Sailing Program (20YF1422100).

## Author Contributions

Zhongbin Su, Conceptualization, Data curation, Formal analysis, Validation, Investigation, Visualization, Methodology, Writing – original draft; Xiaolin Zhou, Supervision, Funding acquisition, Writing – review and editing; Lihui Wang, Conceptualization, Validation, Supervision, Funding acquisition, Writing – original draft, Writing – review and editing.

## Data Availability

Data and codes have been deposited at OSF, accession code: osf.io/h5dwk/.

## Supplementary Materials

### Fast Fourier transform (FFT) analysis of epoched data

To show the neural entrainment by the rhythmic visual stimuli, a data-driven spectral analysis was conducted on the pre-target activity over the visual cortex using a fast-Fourier transform (FFT). For each of the occipital channels (Oz, O1, O2, POz, PO3, PO4, PO7, PO8), epoched data with a time range of 5 cycles relative to the target onset was included for FFT analysis, rendering a time range of −4000 to 0 ms for S+P- (1.25 Hz * 5 cycles), and a time range of - 2000 to 0 ms for S+P+ and S-P+ (2.5 Hz * 5 cycles). FFT analysis was then applied to the frequencies ranging from 0.25Hz to 250Hz, in steps of 0.25Hz. The zero-padding method was used to improve frequency resolution. For each condition, the obtained spectral amplitudes were averaged across all epochs and channels. For each of the three positions (S+P-, S+P+, S-P+), the amplitude difference between regular and irregular conditions was compared with *Paired-T* tests (one-tailed). FDR correction was applied for multiple comparisons across 0-5 Hz frequencies.

**Figure S1.**
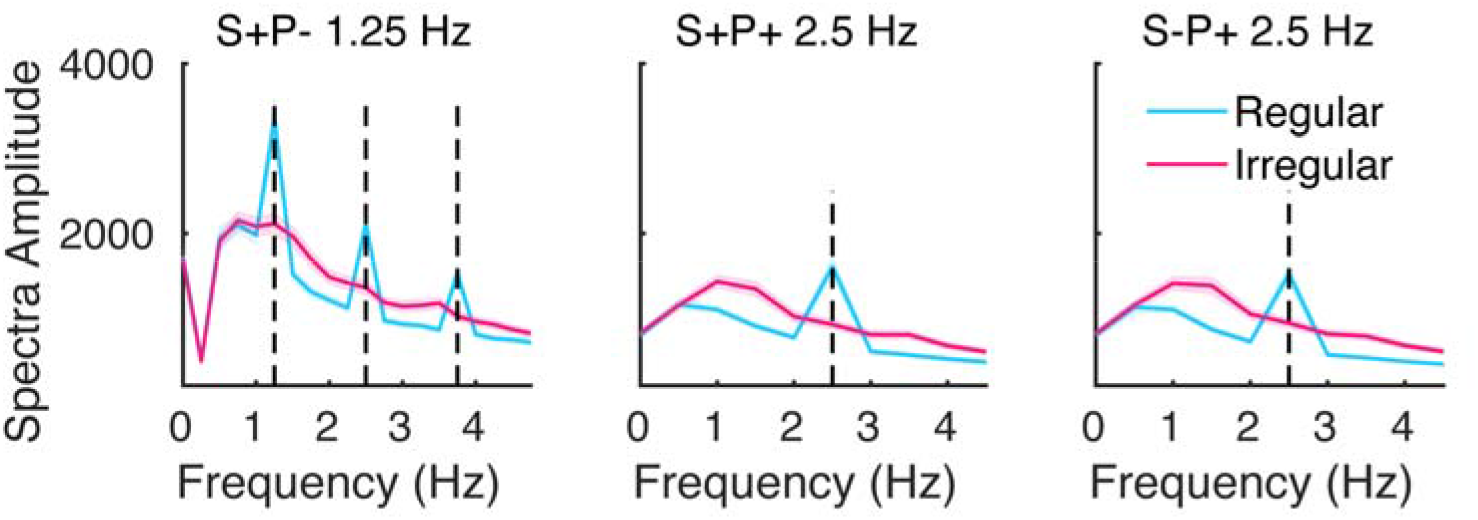
Spectrum amplitudes of the pre-target data over the occipital cortex are shown as a function of frequency for each condition. The neural data was entrained by periodic external stimuli. S+P-, 1.25Hz; S+P+, 2.5Hz; S-P+, 2.5Hz. The shadows indicate SEM across participants.

At S+P- (Figure S1, left), the amplitudes of 1.25Hz and its harmonic frequency (2.5Hz, 3.75 Hz) were higher in the regular condition than in the irregular condition (*paired-T* test, FDR corrected *p* < 0.05). At S+P+ and S-P+ (Figure S1, middle and right), the amplitudes of 2.5Hz were higher in the regular condition than in the irregular condition *(paired-T* test, FDR corrected *p* < 0.05).

### Time-frequency analysis

For each participant, each electrode, and each epoch, a time-frequency transformation was performed on the interval of −4000 to 1500ms relative to the target onset by convolving the data using a Morlet wavelet *(newtimef,* 2021 version EEGLAB). The long epochs were selected to reduce the loss of low-frequency information. The wavelet size was changed linearly from 3 cycles to 25 cycles with frequency from 3 Hz to 50 Hz, in a step of 1Hz. The amplitude spectra were extracted from the complexes and log-transformed. The frequency amplitudes were averaged over electrodes and epochs and then compared between regular and irregular conditions for each time-frequency point with *paired-T* test for each position. Cluster-based permutation was used for multiple comparisons across time points and frequencies. The time-frequency results are shown in Figure S2.

**Figure S2.**
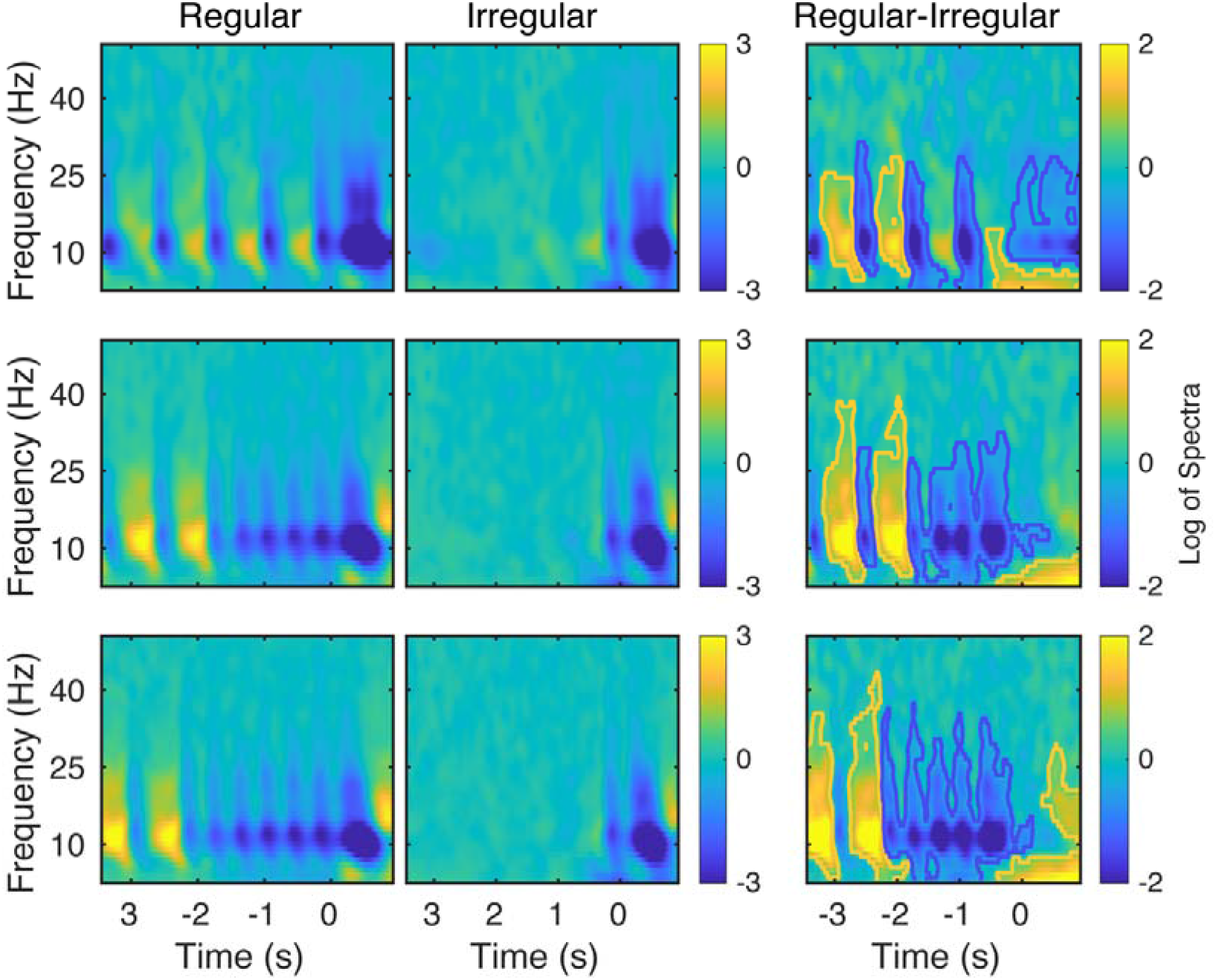
Time-frequency results. The amplitudes are shown as a function of time (relative to target onset) and frequency for each experimental condition (left and middle panels). The difference in time-frequency amplitude between regular and irregular conditions (right panel). The time-frequency clusters marked by the orange and blue lines denote significant differences (orange for positive values and blue for negative values) between regular and irregular conditions *(cluster-based permutation corrected p < 0.05).*

